# Interactome comparison of human embryonic stem cell lines with the inner cell mass and trophectoderm

**DOI:** 10.1101/411439

**Authors:** Adam Stevens, Helen Smith, Terence Garner, Ben Minogue, Sharon Sneddon, Lisa Shaw, Maria Keramari, Rachel Oldershaw, Nicola Bates, Daniel R Brison, Susan J Kimber

**Author notes:** Authors contributed equally to this work (Adam Stevens, Helen Smith). Address all correspondence to: Susan J Kimber, Faculty of Biology Medicine and Health, Michael Smith Building, Oxford Road, Manchester M13 9PT, UK.

## Abstract

Human embryonic stem cells (hESCs) derived from the pluripotent Inner cell mass (ICM) of the blastocyst are fundamental tools for understanding human development, yet are not identical to their tissue of origin. To investigate this divergence we compared the transcriptomes of genetically paired ICM and trophectoderm (TE) samples with three hESC lines: MAN1, HUES3 and HUES7 at similar passage. We generated inferred interactome networks unique to the ICM or TE using transcriptomic data, and defined a hierarchy of modules (highly connected regions with shared function). We compared network properties and the modular hierarchy and show that the three hESC lines had limited overlap with the ICM specific transcriptome (6%-12%). However, this overlap was enriched for network properties related to transcriptional activity in ICM (p=0.016); greatest in MAN1 compared to HUES3 (p=0.048) or HUES7 (p=0.012). The hierarchy of modules in the ICM interactome contained a greater proportion of MAN1 specific gene expression (46%) compared to HUES3 (28%) and HUES7 (25%) (p=9.0×10^−4^).

These findings show that traditional methods based on transcriptome overlap are not sufficient to identify divergence of hESCs from ICM. Our approach also provides a valuable approach to the quantification of differences between hESC lines.

## Introduction

Embryonic stem cell lines are generally derived from the inner cell mass of the preimplantation blastocyst. The proteins OCT4 (*POU5F1*), SOX2 and NANOG are core pluripotency-associated factors that define a network of interactions involved in self-renewal and maintenance of the pluripotent state for human and mouse embryonic stem cells (Boyer, et al., 2005). Each of the core pluripotency factors has been detected in at least some early trophoblast cells, however, they have often not been detected in all cells of the inner cell mass (ICM)/epiblast, for a given embryo (Cauffman, et al., 2009; Kimber, et al., 2008). This heterogeneity has been confirmed by RNAseq analysis of single human preimplantation epiblast cells (Petropoulos, et al., 2016). Recently the central role of OCT4 not only in maintenance of the inner cell mass stem cell population but also in the differentiation of the extra-embryonic trophectoderm (TE) has been established using CRISPR/Cas 9 gene editing in human preimplantation embryos and embryonic stem cells (ESCs)(Fogarty, et al., 2017). Data from the mouse and cynomolgus monkey indicate that the ICM generates a series of epiblast states before giving rise, after implantation, to progenitors of differentiated lineages (Han, et al., 2010; Nakamura, et al., 2016; Weinberger, et al., 2016). At the same time, pluripotency–associated transcriptional networks continue to be expressed in the preimplantation human epiblast (Niakan and Eggan, 2013; Petropoulos, et al., 2016) and early post-implantation cynomolgus epiblast (Nakamura, et al., 2016). Thus, the preimplantation epiblast has transcriptional heterogeneity which is likely to relate to initiation of differentiation events that take place in the early post implantation epiblast and will also impact the generation of ESC lines.

Expression of a number of genes has been associated with the development of extraembryonic cell lineages including *Tead4* (Nishioka, et al., 2008), *Tsfap2c* (Kuckenberg, et al., 2012), *Gata3* (Home, et al., 2009) and *Cdx2* (Strumpf, et al., 2005). But there is evidence suggesting divergence between species in the utilisation of some of these genes such as the Gata family (Grabarek, et al., 2012; Rossant, et al., 2003; Schrode, et al., 2013; Stephenson, et al., 2012) known to play a role in TE generation (Nakamura, et al., 2016). These observations imply that networks of interacting co-regulated proteins might distinguish the transiently pluripotent ICM/preimplantation epiblast from the early differentiated trophectoderm (TE) in a species-specific manner.

In mouse the ground state pluripotency of the ICM appears to be maintained in murine ESCs derived from the ICM and cultured in the presence of LIF together with MEK and GSK3β inhibitors (Weinberger, et al., 2016). This is not the case for human ESCs derived from day 6-7 blastocysts and cultured in standard medium with TGFβ family molecules and FGF-2. It is established in the literature that human ESC lines have more similarities to the murine epiblast after implantation (Faial, et al., 2015; Tesar, et al., 2007) than to the murine ICM and ESCs. In order to understand this difference, it is important to determine how similar hESCs are to the *human* ICM.

Transcriptional analysis of isolated ICM and TE samples from individual human embryos has also been performed, highlighting key metabolic and signalling pathways (Adjaye, et al., 2005). A recent study of 1529 individual cells from 88 human preimplantation embryos defined a transcriptional atlas of this stage of human development (Petropoulos, et al., 2016), however cell lineage allocation can be problematic and inter-individual heterogeneity has been shown to have a major effect on gene expression (Smith, et al., 2019; Stirparo, et al., 2018). Together these data show the relevance of transcriptome based analysis and highlight the need for approaches that account for inter-individual variation.

Recently the heterogeneity present in available human blastocyst single cell RNAseq data has been commented on and sample preparation methods have been questioned (Stirparo, et al., 2018). In the work presented here we have set out to examine how far the gene expression profiles of ICM and TE have diverged from one another at the blastocyst stage, when hESC derivation occurs, and to compare these data to the transcriptome of hESCs using sets of transcriptomic data independent of preparation method. We have defined paired transcriptomic data sets unique to the ICM and TE from the same human embryo. Combined with accepted lists of genes that have differential expression between ICM and TE defined by meta-analysis, we generated ICM-and TE-specific interactome network models. This approach has allowed us to use quantitative network analysis to compare both TE and ICM with hESCs and to evaluate the extent of similarity between ICM/TE and hESC cell lines as well as the hESC lines with each other. These analyses provide an important framework which highlights the development origins of hESCs.

## Results

### Similarities between the transcriptome of inner cell mass, trophectoderm and human embryonic stem cell lines

We used the significant transcriptomic differences between ICM and TE (512 genes) identified by Stirparo *et al* in their meta-analysis of human blastocyst single cell RNAseq data (Stirparo, et al., 2018) to map the relationship of our stem cell transcriptomic data (**Figure 1A**). These data demonstrated that the MAN1, HUES3 and HUES7 transcriptomes identified using frozen RMA (McCall, 2015) were similar to those hESCs previously examined by Yan et al (Yan, et al., 2013) and were in the direction of the NANOG eigenvector. We also observed heterogeneity in the blastocyst single cell RNAseq from Petropoulos *et al* as previously indicated by Stirparo *et al* (Stirparo, et al., 2018).

**Figure 1.**
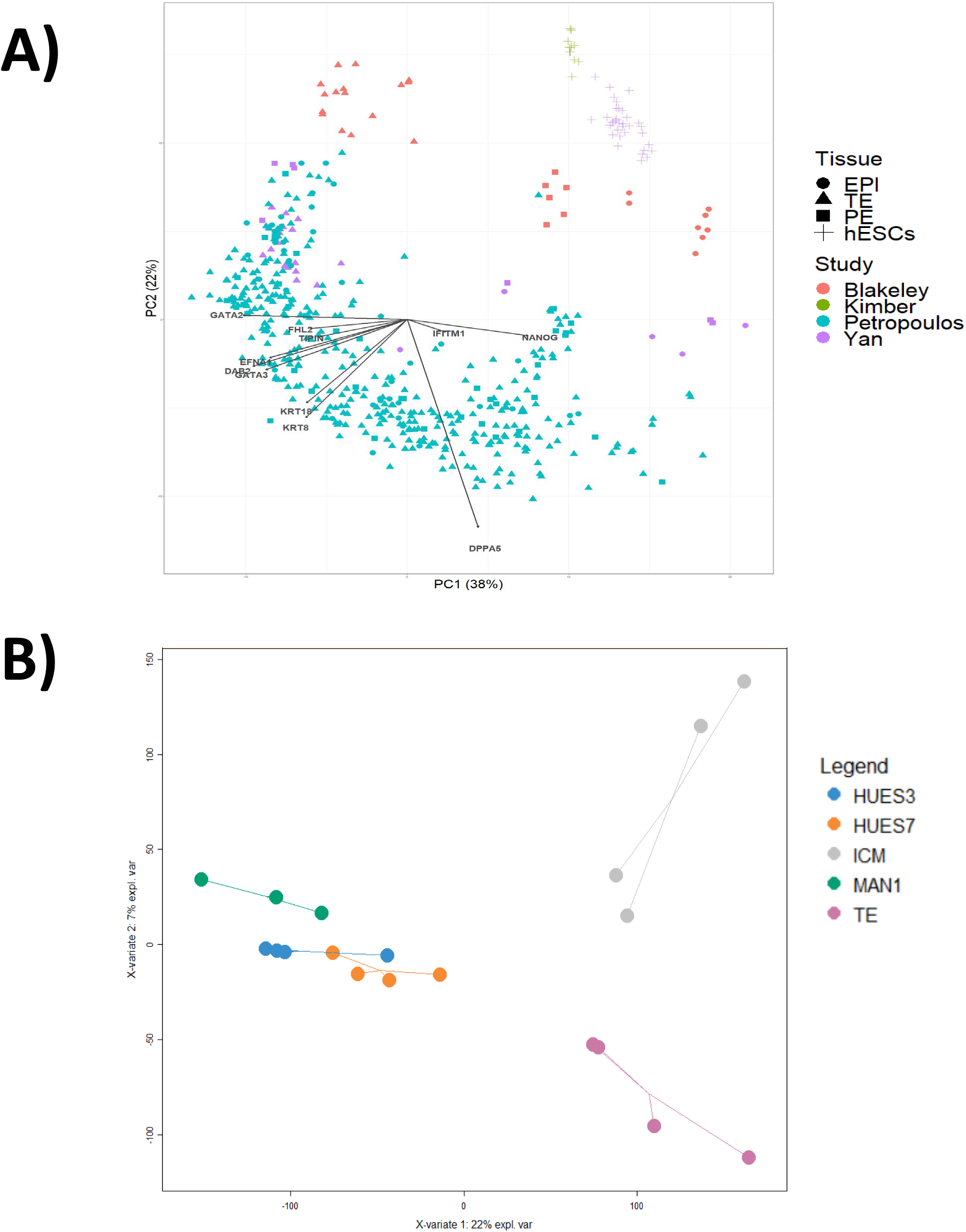
Distance between the transcriptomes of inner cell mass, trophectoderm and human embryonic cell lines as a measure of similarity. **A)** Principal component analysis (PCA) using genes defined by Stirparo *et al* as distinguishing trophectoderm from epiblast. Performed using matching genes across 4 studies, including embryonic tissue and stem cells. Arrows show eigen vectors demonstrating the contributions of key genes to principal components. N-samples=520, N-genes=452. **B)** Gene expression over the entire transcriptome (54613 gene probesets) was defined using the gene barcode approach as a z-score in comparison to a database of 63331 examples of HGU133plus2.0. The Euclidean distances between samples were assessed using partial least square discriminant analysis (PLSDA). Two components are used (X-variate 1 & 2) and the amount of explained variance is listed on the axis. The star plot shows sample distance from the centroid, the arithmetic mean position of all the points in each group.

Frozen RMA barcode Z scores (McCall, 2015; McCall, et al., 2010) for the entire transcriptome (n=54613 gene probe sets) were compared using partial least squares discriminant analysis (PLSDA) to assess the relationship between ICM, TE and the hESC lines MAN1, HUES3, HUES7 (**Figure 1B**). The hESC sample groups were distinct from each other and from ICM and TE (p<0.05). All three hESC cell lines were of equivalent distance from both ICM and TE along the X-axis (X-variate 1), however along the Y-axis (X-variate 2) MAN1 was closer to ICM than HUES3 or HUES7. Similar results were shown with PCA (data not shown).

### Gene expression unique to inner cell mass and trophectoderm and associated gene ontology

Frozen RMA gene barcode was used to isolate gene probe sets present in each embryonic tissue resulting in 2238 probe sets in ICM and 2484 probe sets in TE. These data were used to determine the overlap and unique gene expression in each of these blastocyst tissues (**Figure 2A**). We found 881 and 1227 gene probe sets uniquely expressed in the ICM and TE respectively, corresponding to 719 and 924 unique genes (**Supplemental Table S1**). The genes defined as having unique expression in ICM or TE significantly overlapped with single cell RNA-seq data from human epiblast and trophectoderm cells respectively (both p<1.0×10^−4^), identified in previously published analysis (Petropoulos, et al., 2016). Recognising the potential heterogeneity of samples within the Petropoulos data highlighted by Stirparo et al we used the genes identified by frozen RMA as unique to ICM and TE in combination to categorise the available single cell RNAseq blastocyst data. This analysis resulted in almost perfect classification of the single cell RNAseq datasets from Yan *et al* (Yan, et al., 2013) and Blakely *et al* (Blakeley, et al., 2015) and, as previously shown by Stirparo *et al* (Stirparo, et al., 2018), highlighted the heterogeneity within the Petropoulos data (Petropoulos, et al., 2016) (**Figure 2B**). The stem cell transcriptomic data generated by frozen RMA was no longer proximal to the stem cell data generated by Yan *et al* (Yan, et al., 2013) but had moved further along the eigenvector implying ICM classification (**Figure 2B**).

**Figure 2.**
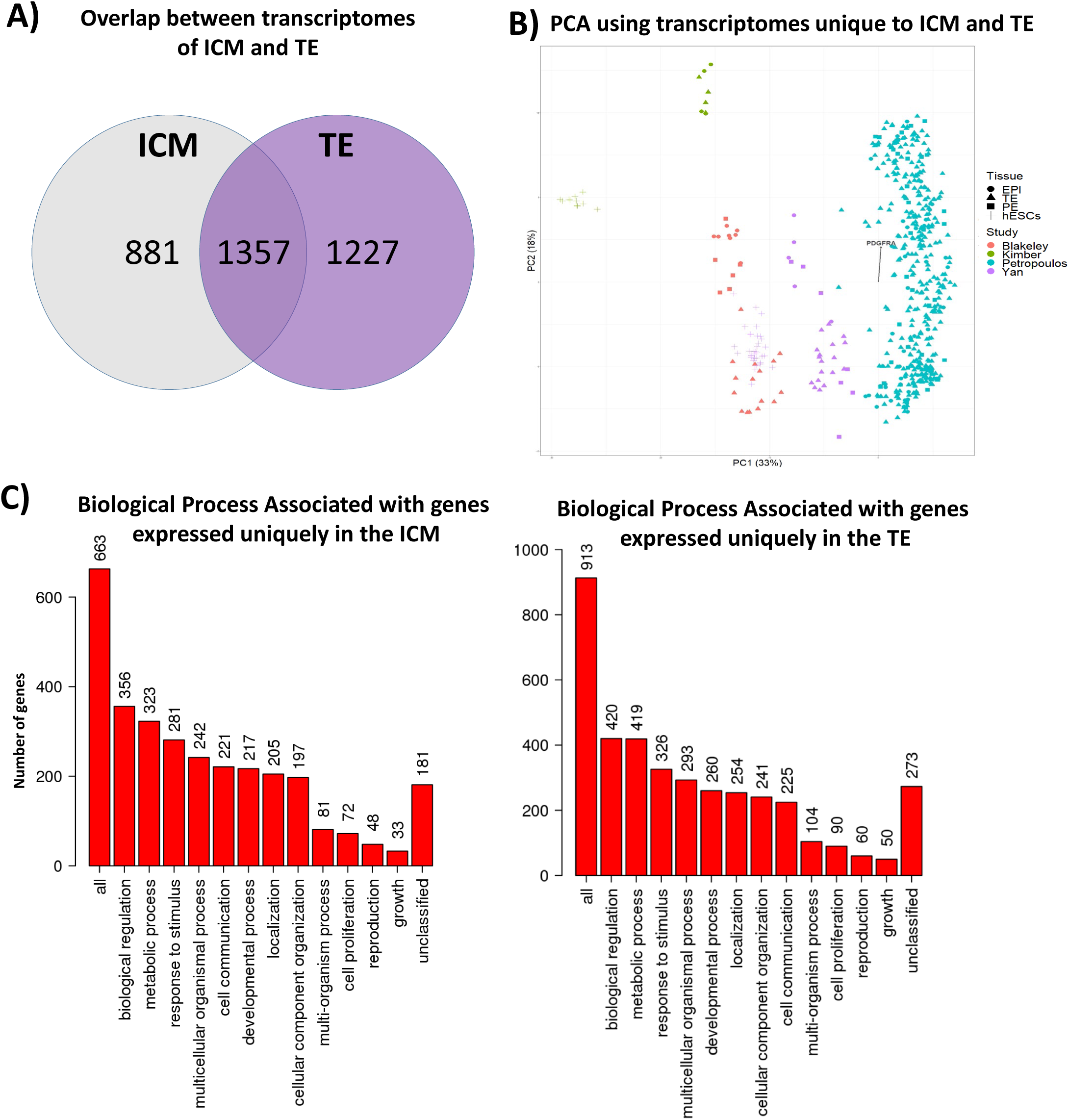
Inner cell mass and trophectoderm specific transcriptome and associated gene ontology. Gene expression over the entire transcriptome was assigned as present or absent using the gene barcode approach, present was defined as a z-score ≥ 5.0 for a gene probeset in comparison to a database of 63331 examples of hgu133plus2.0. This resulted in a set of 2238 gene probesets in ICM and 2484 gene probesets in TE. **A)** A Venn diagram showing the overlap and unique expression of gene probesets in the ICM and TE. **B)** PCA using genes defined to distinguish trophectoderm and epiblast by gene barcode on embryonic cells. N-samples=528, N-genes=452. **C)** Biological process gene ontology (GO Slim) for 663/719 genes used from 881 gene probesets uniquely mapped to the ICM and 913/924 genes used from 1227 gene probesets uniquely mapped to TE.

The genes associated with ICM and TE were grouped by “biological process” ontology showing a similar proportion and ordering in both gene sets, the only difference being a reduction in the proportion of genes of the category “cell communication” in the TE compared to the ICM (**Figure 2C**). More detailed comparison of biological pathways identified “epithelial adherens junction signalling” (ICM p=4.2 x 10^−5^, TE p=7.3 x 10^−4^) as strongly associated with both TE and ICM, and EIF2 translation initiation activity (TE p=4.4×10^−6^, ICM p =0.39) as significantly associated with TE, consistent with the TE being at an early stage of diverging differentiation towards trophectoderm epithelium (Marikawa and Alarcon, 2012), with an active requirement for new biosynthesis (Hasegawa, et al., 2015) (**Supplemental Table S2**).

It was noted that NANOG regulation was strongly associated with the ICM (p=5.9×10^−6^) but not the TE and that CDX2 regulation was associated with TE (p= 9.8×10^−3^) but not ICM, as would be anticipated (Niakan and Eggan, 2013). Using causal network analysis we identified master regulators of gene expression associated with the transcriptomic data. This approach identified MYC (p=7.6×10^−8^), a co-ordinator of OCT4 activity (Fang, et al., 2016), and ONECUT1 (HNF6) (p=4.0×10^−8^), a regulator of the development of epithelial cells (Pierreux, et al., 2006), as the most significantly associated regulatory factors in ICM and TE respectively (**Supplemental Tables S3**).

### Similarities between the inner cell mass and trophectoderm unique transcriptomes and the transcriptome of human embryonic stem cells

Firstly we used the 512 genes, defined by meta-analysis (Stirparo, et al., 2018), as differentially expressed between ICM and TE to quantify correlations with gene expression within the transcriptomes of the hESC lines using hypernetwork analysis. These data highlighted MAN1 as having quantifiably more correlations (1.8 fold p< 1×10^−5^) compared to HUES3 or HUES7 with gene expression that is associated with the differentiation of ICM and TE (**Figure 3A**). The rank order of the stem cell lines was MAN1 >> HUES3 > HUES7 as indicated by the number of co-expressed genes in the interactome (increased proportion of yellow in the heatmap - **Figure 3A**).

**Figure 3.**
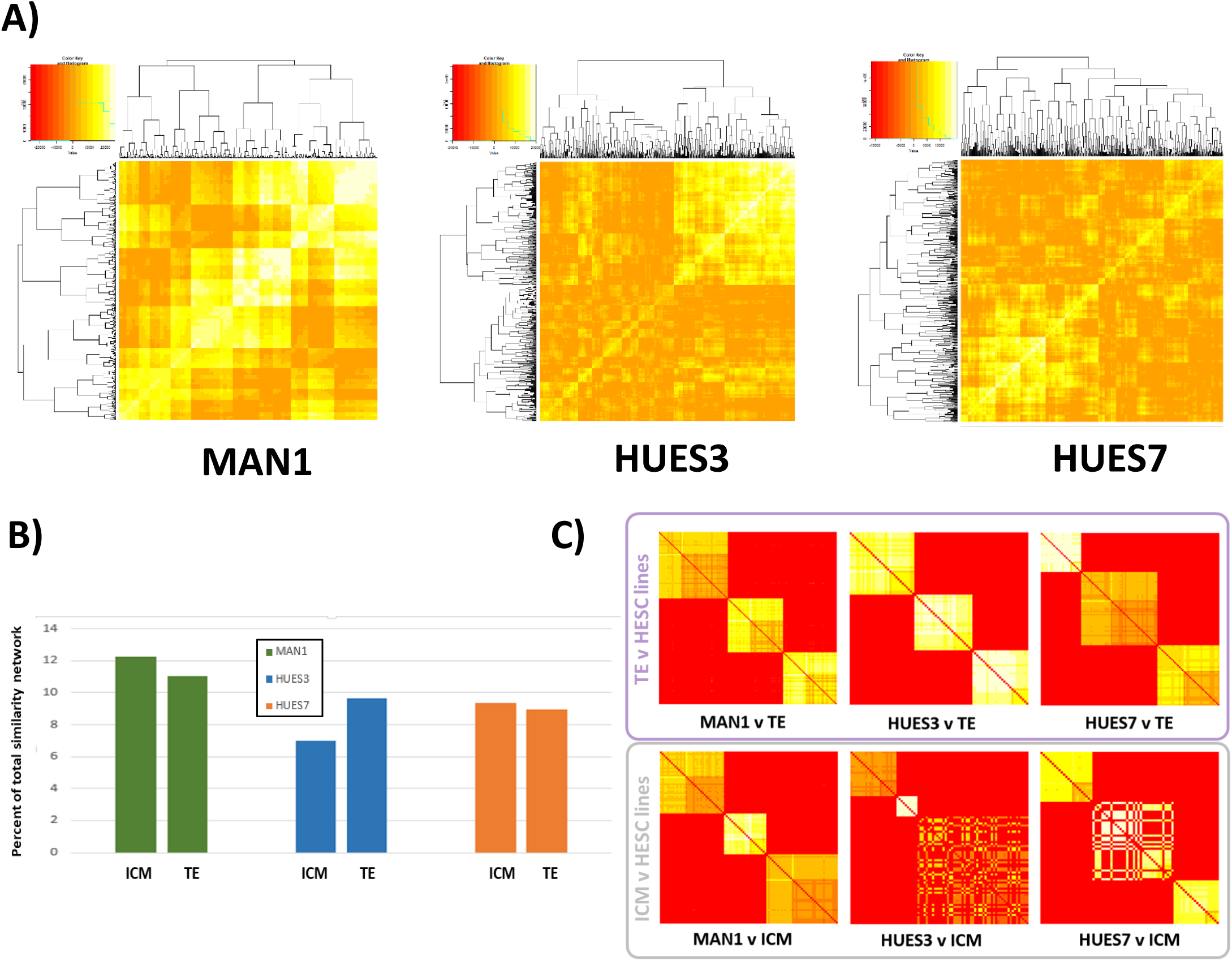
Similarity network fusion to compare homology between the transcriptome of inner cell mass and trophectoderm and human embryonic stem cell lines. Similarity network fusion matrix showing similarity groups between the uniquely expressed ICM and TE gene probesets and the human embryonic stem cell lines (square matrix of gene probesets with leading diagonal showing equivalence mapped to red). Similarity is coloured by intensity from white to yellow, red is dissimilar. Groups of genes with similar expression patterns across both comparisons appear as yellow, whilst those with dissimilar patterns of expression within or between cell lines appear red. Clusters therefore represent genes whose expression patterns are similar to one another both within and between input datasets. Similarity measures not only distance between ICM and the human embryonic stem cell lines but also coherency based on 15 nearest neighbours. Hypernetworks of genes distinguishing trophectoderm from inner cell mass correlated against all other genes in three stem cell lines (HUES3, HUES7, MAN1). Yellow denotes high connectivity between epiblast-distinguishing genes and others. **A)** Proportion of gene probesets in ICM or TE that are similar to human embryonic cell line transcriptome (**Supplementary Figure S1**). **B)** Similarity groups between ICM or TE and the human embryonic stem cell lines forming three clusters. Coherency in gene expression patterns with nearest neighbours is indicated by uniform yellow intensity.

Similarity Network Fusion (SNF) was used to assess the similarity of gene expression patterns between cell lines and ICM or TE. SNF uses nearest neighbour component to its algorithm to identify regions where this pattern is *coherent*. A region of coherency across a stem cell line and either TE or ICM represents a group of genes whose expression pattern is conserved between embryonic tissue and hESCs. The analysis highlighted a limited similarity of hESC lines with ICM (between 6% and 12% similarity) and TE (between 9% and 11%), consistent with the distance between the hESC lines and TE and ICM as observed by PLSDA analysis (**Figure 3B & Supplemental Figure S1**). Three primary clusters of similarity were identified in all comparisons between the hESC lines and ICM or TE (**Figure 3C**). These clusters were of equivalent similarity in TE with all hESC lines, as indicated by uniform yellow intensity indicating coherency with nearest co-expressed neighbours, implying highly co-ordinated expression. However, when ICM was compared with hESCs, coherency was noted only with MAN1 and not with the other hESC lines (**Figure 3C**).

### An interactome network model of gene expression unique to ICM can be used as a framework to assess similarity with human embryonic stem cells

An interactome network model can be used to consider the proteins derived from differentially expressed genes and the proteins that they interact with. Using this approach allowed us to consider the wider biological interactions generated by the gene expression unique to either the ICM or TE and to implement these models as a framework to assess similarity with the hESC lines.

We used the genes with differential expression between ICM and TE as defined by Stirparo *et al* (Stirparo, et al., 2018) to generate ICM and TE specific network models by using the genes with positive fold change in expression in each specific tissue (337 for ICM and 175 for TE) as a basis for network inference. We also separately used the genes with unique expression in either ICM or TE (**Figure 2A**) to generate interactome network models by inference to known protein-protein interactions (**Supplemental Figure S2A & 2B**).

As interactome networks account for inferred interactions these may be shared between different network models. Comparing the TE and ICM interactome network models an overlap of 2517 and 5659 inferred genes was present in the Stirparo models and the models based on our de novo data respectively. These overlaps represent protein:protein interactions, accounting for 85% (Stirparo) and 72% (our model) of the ICM interactome along with 66% (Stirparo) and 30% (our model) of the TE interactome.

We examined further the network models based on the uniquely expressed genes in ICM and TE identified by frozen RMA. Both networks were enriched for genes associated with pluripotency, for example NANOG with the ICM network and CDX2 within the TE network, as identified by gene ontology analysis. The ICM network contained 93/167 and 161/240 genes and the TE network contained 94/167 and 185/240 genes related to core pluripotency associated factors by RNAi (Ding, et al., 2009; Hu, et al., 2009; Ivanova, et al., 2006; Ng and Lufkin, 2011; Zhang, et al., 2006) and protein interaction (Liang, et al., 2008; Ng and Lufkin, 2011; Pardo, et al., 2010; van den Berg, et al., 2010; Wang, et al., 2006) screens respectively. The similarity of TE with ICM networks for pluripotency factors is likely to reflect the fact that this tissue has only very recently begun to diverge.

Using uniquely expressed genes derived from our de novo transcriptomic analysis we were able to determine the shared transcriptome between ICM or TE and each human embryonic stem cell line and map these onto the respective ICM or TE interactome network model. Of the genes shared between the hESC lines and ICM there was no difference between the proportions each line shared with the network model (p=0.74), for the genes shared between the hESC lines and TE, MAN1 had a significantly smaller proportion of genes shared with the TE network model (p= 0.03).

### Similarities and differences in topology between human embryonic stem cell lines in relation to inner cell mass and trophectoderm network models

As the ICM and TE interactome models shared a significant proportion of the same genes, we went on to assess the network topology of these models to determine further similarities and differences with the genes shared with the hESC lines. Analysis of the network topology of the ICM and TE interactome demonstrated that the genes shared with the hESC lines were enriched for highly connected genes (as measured by degree, the number of interactions made to other genes). We found that HUES3 and MAN1 were more connected than HUES7 in the ICM network (MAN1vsHUES7 p=0.04, HUES3vsHUES7 p=0.04, MAN1vsHUES3 p=0.94) but not in the TE network (MAN1vsHUES7 p=0.21, HUES3vsHUES7 p=0.28, MAN1vsHUES3 p=0.89) (**Figure 4A & 4B**).

**Figure 4.**
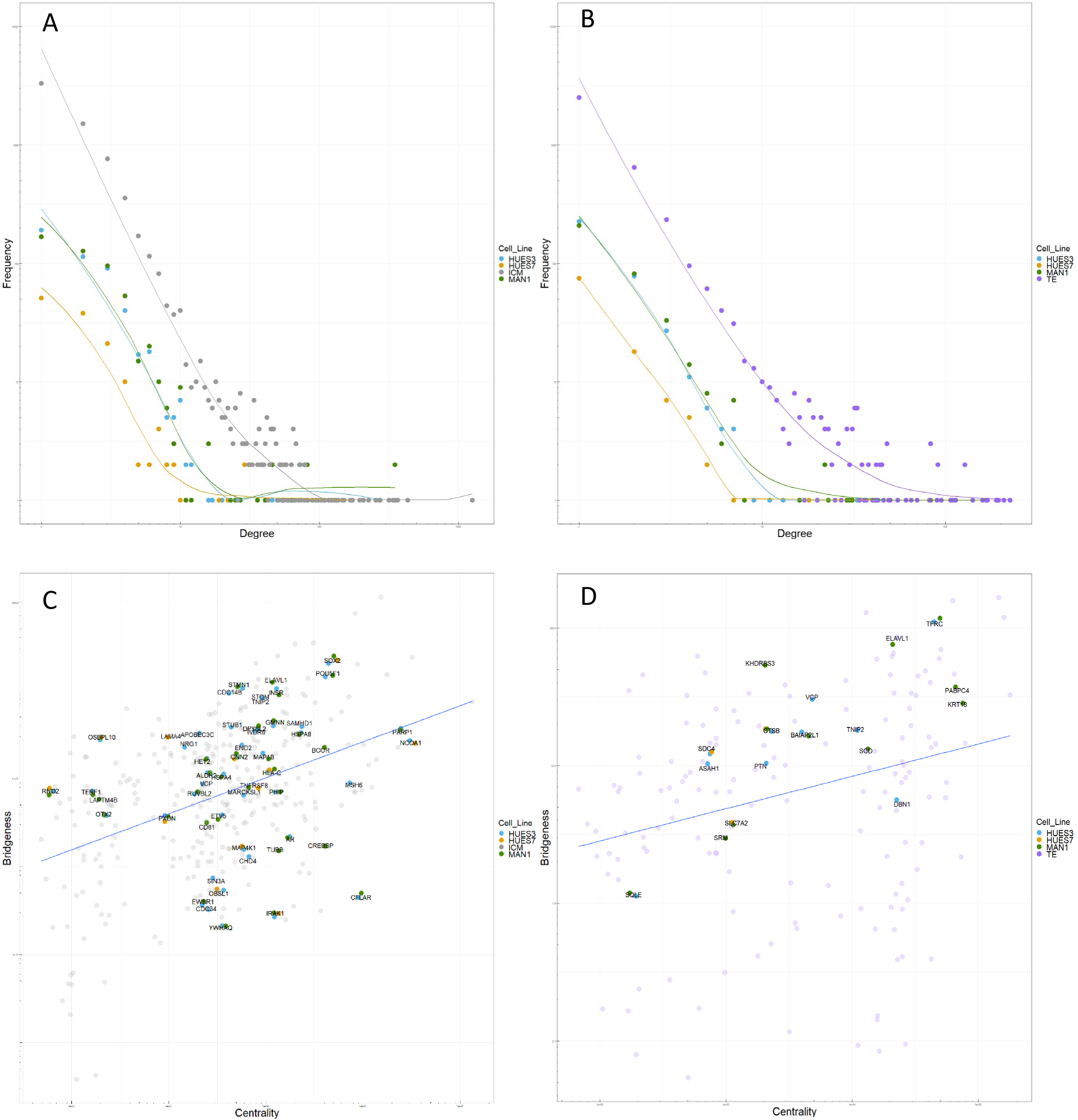
The network topology of the ICM interactome is enriched in human embryonic stem cells. Degree distribution of unique genes in an inner cell mass (**A**) and trophectoderm (**B**) network model. HUES3, HUES7 and MAN1 are subsets of TE or ICM. These plots demonstrate that HUES3 and MAN1 are more connected than HUES7 in both networks. Plots are of log frequency and log degree. Bridgeness vs centrality measures in an inner cell mass (**C**) and trophectoderm (**D**) network model. HUES3,HUES7 and MAN1 are subsets of TE or ICM. HUES3 and MAN1 have a greater proportion of date-like hubs than HUES7 or either ICM or TE, demonstrating an increased number of genes with network properties of transcription factors.

To further investigate the putative functional relevance of genes shared between the ICM or TE interactome models and the hESC lines we determined whether these genes had “party” or “date” like properties. In protein interaction networks party hubs co-ordinate local activity by protein complexes, whereas date hubs regulate global effects and are assumed to represent the transient interactions that occur with transcription factors (Agarwal, et al., 2010; Chang, et al., 2013). Date-like network hubs have been shown to possess a higher “bridgeness” property at any position within the interactome (Kovacs, et al., 2010). Bridgeness is a network property that measures overlap between network modules and this score can be compared at different positions within the network by plotting it against “centrality”, a network property that measures the influence of a node in a network (Kovacs, et al., 2010). HUES3 and MAN1 were shown to have a greater proportion of date-like hubs than HUES7 in either the ICM or TE network models based on the Stirparo data, demonstrating an increased number of genes with network properties of transcription factors. (**Figure 4C & 4D**). This observation implies an enrichment for date-like network hubs in the genes shared between the hESC lines and the ICM or TE interactome network models, implying in turn an enrichment of transcription factor activity. Using the interactome models derived from de novo transcriptomic analysis the network topology of the ICM and TE demonstrated that the genes shared with the hESC lines were enriched for highly connected genes (as measured by degree, the number of interactions made to other genes) and the enrichment seen was not statistically different between the hESC lines (**Figure 5A & 5B**).

**Figure 5.**
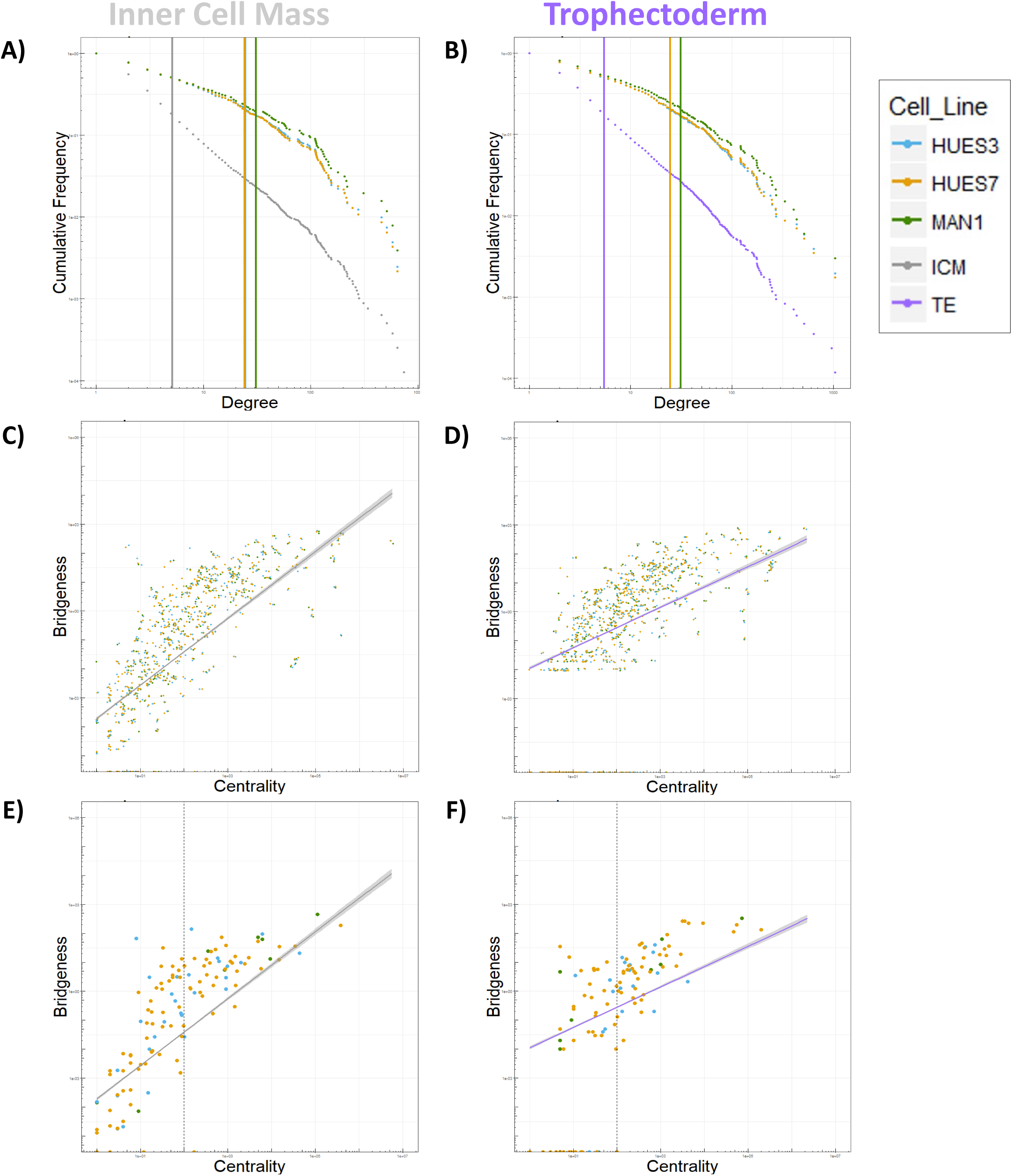
The network topology of the ICM interactome based on the unique expression of genes compared to TE is enriched in human embryonic stem cells. **A)** ICM interactome connectivity and **B)** TE interactome network connectivity as measured by the degree (connectivity) of each gene within the network model (x-axis) plotted against the frequency of that connectivity within the network (y-axis). Plots are of log frequency and log degree. **C)** ICM interactome and **D)** TE interactome centrality score (x-axis), a network property that measures the influence of a node, plotted against bridgeness (y-axis), a network property measuring the bridge-like role of genes between network modules. The line with 95% confidence intervals shaded represents the centrality and bridgeness values over the entire network, genes shared with the human embryonic stem cells are marked. **E)** ICM interactome and **F)** TE interactome centrality versus bridgeness shown for genes uniquely expressed in each human embryonic stem cell line. Dotted vertical line placed at centrality value of 100 separates two perceived trajectories in the data.

All three hESC lines were shown to be enriched for bridgeness score in relation to centrality when compared to the full ICM or TE networks based on de novo transcriptomic data (**Figure 5C & 5D**). We identified the overlap of genes expressed in the ICM or TE and the hESC cell lines (**Supplemental Figure S2**). There were 590 and 652 shared genes between all the three hESC lines and ICM or TE respectively (**Supplemental Figure S3A & S3B**). When we examined genes uniquely expressed in each of the hESC lines (**Supplemental Figure S3A & S3B**), the highly central genes in both networks (centrality score >100) were significantly enriched for bridgeness in ICM (p=0.016) but not TE (p=0.105), indicating more date-like properties in ICM (**Figure 5E & 5F**). In the ICM interactome network model MAN1 was significantly more date-like than HUES3 (p=0.048) and HUES7 (p=0.012). This observation implies that the MAN1 cell line shared significantly more transcription factor activity with ICM which is hierarchically more important within the ICM interactome, than do either HUES3 or HUES7. Biological pathways associated with genes uniquely expressed in each of the hESC lines are shown in **Supplemental Figure S3B**. In MAN1 “PDGF signalling” and “cell cycle control of chromosome replication” were associated with the unique gene expression shared with ICM. PDGF signalling is required for primitive endoderm cell survival in the inner cell mass of the mouse blastocyst (Artus, et al., 2013) and the pluripotency associated transcription factor NANOG (referred to above) has been shown to influence replication timing in the cell cycle (Apostolou, et al., 2013; Hiratani, et al., 2010).

### Modular hierarchy of the ICM and TE interactome network models reveal an enrichment in MAN1 for ICM and an enrichment in HUES7 for TE

Network modules are sub-structures of a network that have a greater number of internal connections than expected by chance. Modules are known to represent functionally related elements of a network and can be ranked hierarchically by their centrality within a network, with the assumption that the more central modules are functionally dominant within the network. We defined modules within our TE and ICM interactome network models allowing for overlap and arranged these into a hierarchy of influence by centrality score (Kovacs, et al., 2010) (**Figure 6A**). The ICM and TE interactome network models had a hierarchy of 109 and 163 along with 71 and 201 modules of different sizes in the models based on the Stirparo data and the de novo data respectively. There was no difference in the proportion of modules compared to network size between the ICM and TE interactome network models (p<0.2) using either the Stirparo data or our own data (**Supplemental Figure S4 & Supplemental Tables S4 & S5**). The robustness of the definition of network modules in the ICM and TE interactome network models based on our de novo transcriptomic data was confirmed by permutation analysis of the proportional random removal of genes (Reimand, 2013) (**Supplemental Figure S5**). This established that the majority of modules were robust to the removal of large proportions of the network, with only 2 of the top 47 ICM and 8 of the top 49 TE modules analysed experiencing a significant (p<0.05) reduction in connectivity within the module following the removal of a random 20% of the network iterated 100 times.

**Figure 6.**
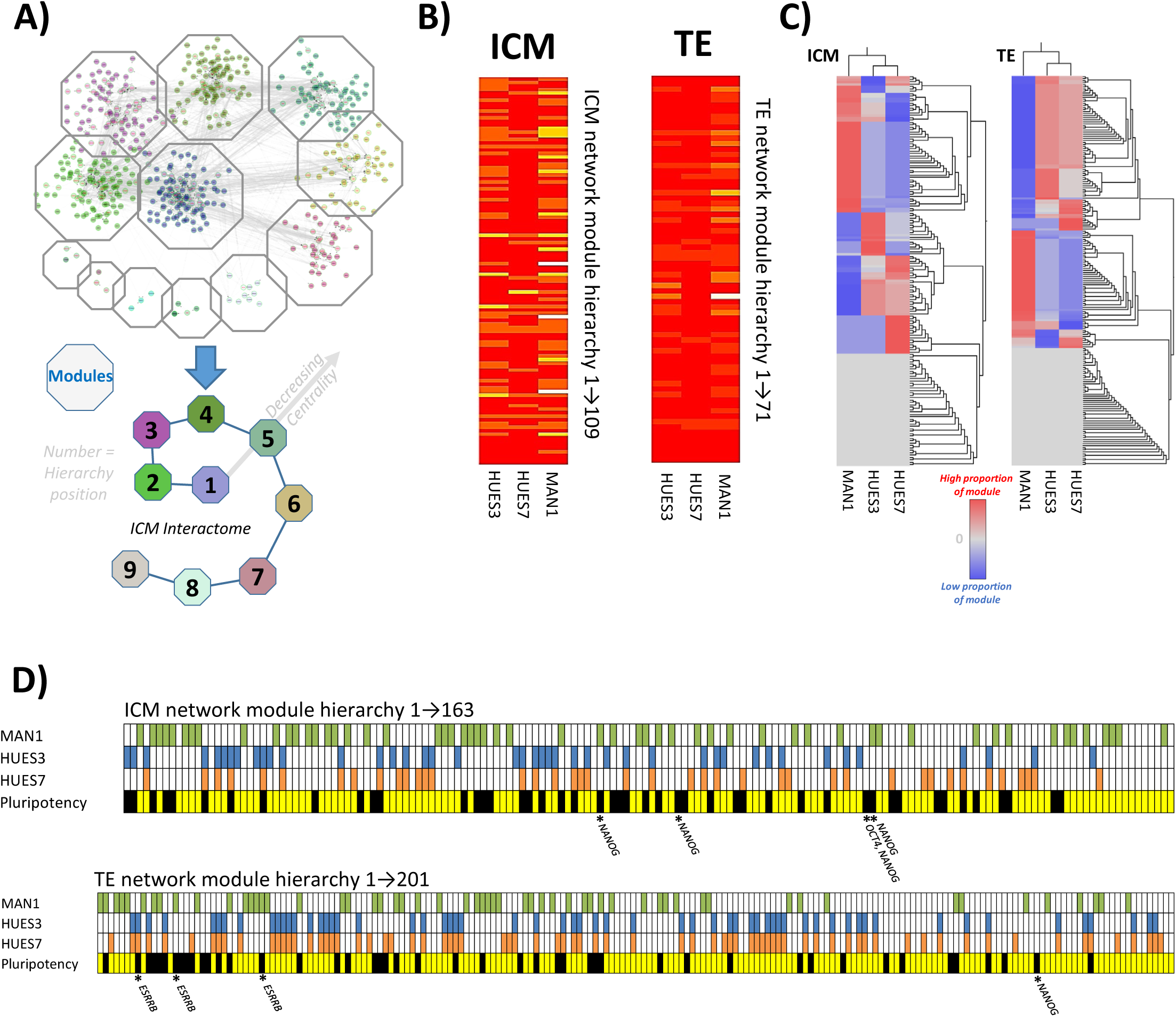
The modular structure of the interactome network model of gene expression unique to ICM and TE can be used as a framework to assess similarity with human embryonic stem cells. A) The modular structure of the ICM interactome was defined using the Moduland algorithm to assess the presence of highly connected gene modules. These were then formed into a hierarchy based on their centrality score, a measurement of network topology related to the influence of a network element on the rest of the network. B) Network module hierarchy in an inner cell mass and trophectoderm network models based on the Stirparo data. Yellow and orange bands demonstrate the presence of genes determined to be unique to each cell line using a gene barcode approach. The greater the intensity of the yellowness, the larger the number of unique genes represented in the meta-node (10 most central nodes in each module). This shows that MAN1 has a greater number of unique genes represented in both TE and ICM networks, particularly in the most central modules and to a greater extent in ICM than TE. C) The proportion of each module shared with the human embryonic stem cell lines was defined and clusters of modules with similar shared gene expression were assessed using a heatmap. D) The clusters of modules with similar proportions shared with specific human embryonic stem cell lines is represented in hierarchical order. Clusters are coloured to mark for which human embryonic cell line they are enriched. Pluripotency track represents which modules contain known pluripotency associated genes in black. An asterisk is used to mark where NANOG, OCT4 and ESRRB are situated in the modular hierarchy.

Network module hierarchy in inner cell mass and trophectoderm network models based on the Stirparo data was assessed and the proportion of each module that also mapped to genes within the transcriptome of the human embryonic stem cell lines. This analysis again showed that MAN1 had a greater number of unique genes represented in both TE (MAN1vsHUES3 p=0.004, MAN1vsHUES7 p=2.95×10^−6^) and ICM (MAN1vsHUES3 p=0.026, MAN1vsHUES7 p=2.95×10^−8^) networks, particularly in the most central modules and to a greater extent in ICM than TE (**Figure 6B**).

The genes with shared expression between ICM or TE networks based on the de novo transcriptomic data and the hESC lines were mapped to each interactome module. In the ICM network 116/163 modules (71%) were still enriched for gene expression shared between hESC lines and ICM. A greater proportion of hESC associated modules in the ICM interactome network model were enriched for MAN1 gene expression (0.46) compared to HUES3 (0.28) and HUES7 (0.25) (p=9.0×10^−4^, chi squared test). In the TE interactome network model 132/201 modules (65%) were enriched for gene expression shared between hESC lines and TE. The smallest proportion of enriched hESC associated modules occurred in HUES7 (0.17) compared to MAN1 (0.39) and HUES3 (0.44) (p=3.1×10^−6^, chi squared test) (**Figure 6C**).

The modules assessed as having enriched gene expression in specific hESC lines were mapped to the module hierarchy in the ICM or TE interactome network model based on the de novo transcriptomic data (**Figure 6D**). These data show an enrichment of the modules that have the greatest proportion of shared gene expression with MAN1 in the upper part of the module hierarchy in both ICM and TE indicating that the MAN1 associated modules were likely to be more functionally active in both the ICM and TE interactomes.

Gene expression uniquely present in each of the hESC lines (**Supplemental Figure S2**) was mapped to the central core (most central 10 genes) of each of the modules in the ICM and TE interactome network models (**Supplemental Figure S3**). This analysis highlighted only gene expression present uniquely in MAN1 or HUES7 in the upper part of the module hierarchy in the ICM and TE interactome network models indicating that HUES3 associated modules had a reduced presence in the function of the ICM. The upper part of the TE network model module hierarchy was enriched for both HUES7 and MAN1 uniquely expressed genes, indicating a dominant effect of these hESC lines on TE function, compared to HUES3.

Finally, relating these analyses to the enrichment for pluripotency associated genes we defined in the ICM and TE interactome models, we examined this relationship to the modular hierarchy of the ICM and TE interactome network models. We assessed whether any of the pluripotent genes mapped to the central core of 10 genes in a network module (coloured black in **Figure 6D**). In the ICM modular hierarchy 16, 13 and 11 of the modules enriched in MAN1, HUES3 and HUES7 respectively also mapped to pluripotency genes. In the TE modular hierarchy 18, 11 and 15 of the modules enriched in MAN1, HUES3 and HUES7 respectively also mapped to pluripotency genes. It was noted that OCT4 (*POU5F1*), a primary marker of ICM (Hochedlinger and Jaenisch, 2015), was present in the central core of the modules from the ICM but not the TE network models. *NANOG*, another marker of ICM (Hochedlinger and Jaenisch, 2015), was present four times in the ICM and only once in the TE network models. Also estrogen-related-receptor beta (*ESRRB)*, a marker of TE (Latos, et al., 2015; Nicola, et al., 2018), was present three times in the TE but not at all in the ICM network models. In the ICM network model, 2 of the 3 NANOG associated modules are enriched for MAN1 gene expression and the module associated with both NANOG and OCT4 had equivalent enrichment in MAN1, HUES3 and HUES7. In the TE network model the NANOG associated module was low in the hierarchy (76/201) and had equivalent enrichment in MAN1, HUES3 and HUES7. In the TE network model the three ESRRB associated modules were at the upper end of the module hierarchy with the highest ranked (8/201) being enriched in HUES3 and HUES7 and the other two being associated with MAN1 (**Figure 5C**). These data combined show that the key transcription factors (and partners) known to be associated with ICM and TE have biologically logical but different associations with hESC lines within the modular hierarchies of the interactome network models.

## Discussion

The analysis presented in this manuscript has defined gene interactome network models of ICM and TE and used these to quantitatively assess the relationship to pluripotency of several human embryonic stem cell lines derived from the ICM.

The MAN1 human embryonic stem cell line was furthest from both ICM and TE using distance metrics on the unsupervised transcriptome. Only ∼10% of genes uniquely expressed by the ICM (compared to TE) were shown to have similarity to expression patterns in MAN1, HUES3 and HUES7 using SNF. However MAN1 was found to be most similar to ICM as it had both a greater enrichment of genes and a greater coherency with nearest neighbours in comparison to HUES3 and HUES7. Substantial enrichment of human embryonic stem cell line gene expression was also observed in relation to TE but, whilst this was shown to be coherent with nearest neighbours, MAN1 and HUES7 showed a reduced similarity compared to that for ICM while HUES3 had an increased similarity to TE.

We used interactome network models of ICM and TE as frameworks to map overlapping gene expression from MAN1, HUES3 and HUES7. Using network topology as a marker of functionality we demonstrated that all the human embryonic stem cell lines had gene interaction networks with increased connectivity in both the ICM and TE interactome network models generated from gene expression data. All human embryonic stem cell lines also showed an enrichment for network topology that was associated more with date hubs than with party hubs, in ICM and TE network models. Date hubs are network positions that are associated with non-concurrent signalling and are more likely to represent transcription factor activity related to the execution of a developmental programme (Agarwal, et al., 2010; Chang, et al., 2013; Kovacs, et al., 2010; Ng and Lufkin, 2011). A key finding of this study is that date hubs central to the network model, and therefore likely to influence a greater proportion of network function, were significantly enriched in the overlap of genes uniquely shared between MAN1 and the ICM compared to genes uniquely shared between HUES3 or HUES7 and ICM.

We defined a functional hierarchy of overlapping network modules in both the ICM and TE interactome network models and used this as a framework to study the relationship of MAN1, HUES3 and HUES7 with ICM and TE gene expression. MAN1 had greater enrichment in the upper hierarchy for both ICM and TE network models both overall and for uniquely expressed genes.

Taken together these observations demonstrate the utility of network approaches to quantify underlying similarities based on the position of transcriptomic differences in an interactome network model. Quantitative comparison of the hierarchy of the ICM and TE interactome network modules in relation to the expressed genes in the human embryonic stem cell lines provided further insight into similarities and differences between the cell lines beyond those defined by traditional distance metrics.

An assessment of master regulators of transcription associated with the ICM and TE specific gene expression identified known tissue specific transcriptional regulators – NANOG in ICM (Hochedlinger and Jaenisch, 2015; Ng and Lufkin, 2011) and CDX2 in TE (Niakan and Eggan, 2013; Niwa, et al., 2005). Both the ICM and TE network models were enriched for genes associated with pluripotency (Hochedlinger and Jaenisch, 2015; Ng and Lufkin, 2011) an observation in alignment with recent diversification of these tissues. The upper part of the hierarchy of network modules in both the ICM and the TE interactome network models was enriched for pluripotency associated genes. However MAN1 was more closely associated with gene modules including NANOG in the ICM interactome network model compared to HUES3 and HUES7 cell lines. In the TE interactome network model HUES3 and HUES7 were associated with the *ESRRB* related module at the highest position in the module hierarchy whilst MAN1 was also primarily associated with two further *ESSRB* related modules. ESSRβ, a direct target of Nanog (Festuccia, et al., 2012), has been shown to be important in murine ES cells as a co-regulator of Oct4 with Nanog (Zhang, et al., 2008) and a regulator of Gata6 though promoter binding (Uranishi, et al., 2016). Using chromosome conformation capture sequencing Nanog interacting modules were found to be more enriched with target sites for Esrrb as well as KLf4, Sox2 and cMyc target sequences with less consistency in Nanog and Oct4 target sequences (Apostolou, et al., 2013). ESSRβ works with p300 to maintain pluripotency networks, generating a permissive chromatin state for binding of Oct4, Nanog and Sox2 and has been implicated in reprogramming epistem cells to an iPSC state (Adachi, et al., 2018).

Overall these data reveal that MAN1 had the greatest similarity to ICM compared to the other hESC lines despite being least related to ICM in the PLSDA analysis. This observation is based on **I)** greater coexpression with other tissue specific gene expression in the hypernetwork analysis, **II)** coherency in the SNF analysis with nearest neighbour genes, **III)** significantly increased proportion of genes with a date-like hub property in the ICM network, **IV)** an increased proportion of genes mapping to ICM interactome network modules and **V)** an association with ICM network gene modules that map to NANOG activity. Concordance has been identified between transcriptomic regulation in human induced pluripotent stem cells and the ICM (Kilens, et al., 2018) but this has not been fully mapped at the level of the interactome. We propose that the network approach presented in this manuscript represents a significant advance on distance metrics in the comparison on hESC lines.

By using a barcode approach to define genes uniquely expressed we were able to define ICM- and TE-specific interactome network models, an important advance from more traditional comparative modelling using differential gene expression (McCall, 2015; McCall, et al., 2014; Zilliox and Irizarry, 2007). We also confirmed similarity of the underlying transcriptomic data with findings from single cell RNAseq data (Petropoulos, et al., 2016) and the independent meta-analysis of that data (Stirparo, et al., 2018) corroborating our observations. These comparisons also confirmed the importance of network structure in the analysis we have undertaken (Rizvi, et al., 2017). We demonstrated the robustness of our network models by establishing module coherency over successive reductions of network model size (by gene removal), therefore establishing a high level of confidence in the analysis of related gene modules and network topology (Reimand, 2013).

The differences between ICM and TE with all three hESC lines may partially reflect the genetic background of the infertile couples donating embryos for analysis and stem cell derivation. Previously we have performed re-analysis of single cell ICM and TE RNAseq from Petropoulos *et al* 2016 (Petropoulos, et al., 2016) and shown a strong effect of inter-individual genetic variation (Smith, et al., 2019). To account for this we have restricted our analysis in this manuscript to only genetically matched pairs of ICM and TE. The similarities we have established by comparison to other work (Petropoulos, et al., 2016) indicate that the data presented in this manuscript is robust to inter-individual differences. The greater dissimilarity of MAN1 to HUES7 and HUES3, revealed in the overlap of the transcriptome to the ICM interactome network modules, may reflect differences in genetic background of individual lines, or derivation regime since HUES3 and HUES7 were derived in the same lab at a similar time (Cowan, et al., 2004; De Sousa, et al., 2009). However it should be noted that all hESC lines were enriched for connectivity, a marker of function, within the ICM interactome, an observation in agreement with a fundamental similarity between hESC lines, despite different genetic background and embryo generation or hESC derivation methods (De Sousa, et al., 2009). It was also noted that hESC lines are different in very many gene modules to ICM. Although the ICMs have totally different genetic background to the hESC lines assessed here, the fact that the hESCs are more dissimilar than the ICMs are to each other does add further weight to this conclusion.

Concern has been raised (Stirparo, et al., 2018) about the heterogeneity of tissue classification in the single cell RNAseq data from Petropoulos *et al* (Petropoulos, et al., 2016). Our work broadly supports these observations but also highlights that tissue classification can be made despite concerns in the sample preparation (Stirparo, et al., 2018) or in inter individual differences (Smith, et al., 2019). This observation would suggest that rigid definitions of tissue specific expression are not necessarily helpful as we expand into single cell analysis. Whilst there is an inherent heterogeneity in the transcriptome of the early embryo that has been defined in the work presented here.

The use of network approaches to quantify similarities between hESCs and their tissue of origin is a developing field. Network summary approaches have been used with promising results (e.g. CellNet (Cahan, et al., 2014)). Correlation networks generated from gene expression have been used to generate quantitative comparison based on the analysis of discrete network modules (Huang, et al., 2014). Network driven approaches can also be used to deal with the large number of comparisons present in the analysis of ‘omic data sets, e.g. topological data analysis (TDA) (Rizvi, et al., 2017) and SNF (Wang, et al., 2014). In the work presented here we have used an efficient method to generate hierarchies of overlapping gene modules (Kovacs, et al., 2010; Szalay-Beko, et al., 2012), thus accounting for the underlying network topology, and supported this analysis using SNF (Wang, et al., 2014) to generate quantitative comparison of hESC lines with ICM and TE. The approach we have developed accounts for both the hierarchy of modules within a network and the large number of comparisons performed in an unsupervised manner to generate robust conclusions. This has allowed us to apply quantitative approaches to determine the similarities of three hESC lines to each other in relation to ICM and TE. We have identified overall similarity of the transcriptomes and we have also defined how these similarities manifest at the level of the interactome. Our findings highlight the diversity inherent in the establishment of hESC lines and also present methods to quantitatively compare similarity and identify key differences using a network approach.

## Methods

### Embryos

Human oocytes and embryos were donated to research with fully informed patient consent and approval from Central Manchester Research Ethics Committee under Human Fertility and Embryology Authority research licences R0026 and R0171. Fresh oocytes and embryos surplus to IVF requirement were obtained from Saint Mary’s Hospital Manchester, graded and prepared as described in Shaw et al 2013 (Shaw, et al., 2013).

### Embryo sample preparation and microarray analysis of transcriptome

Donated embryos were cultured in ISM-1/2 sequential media (Medicult, Jyllinge, Denmark) until blastocyst formation. At embryonic day 6 the zona pellucida of the embryos were removed by brief treatment with Acid Tyrode’s solution pH 5.0 (Sigma-Aldrich, Gillingham, UK), and denuded blastocysts were washed in ISM2 (Medicult). Blastocysts were lysed and reverse transcribed as previously described (Bloor, et al., 2002; Shaw, et al., 2012) and cDNA was prepared by polyA-PCR (Brady and Iscove, 1993) which amplifies all poly-adenylated RNA in a given sample, preserving the relative abundance in the original sample (Al-Taher, et al., 2000; Iscove, et al., 2002). A second round of amplification using EpiAmp™ (Epistem, Manchester, UK) and Biotin-16-dUTP labelling using EpiLabel™ (Epistem) was performed in the Paterson Cancer Research Institute Microarray Facility. For each sample, our minimum inclusion criterion was the expression of β-actin as evaluated by gene-specific PCR. Labelled PolyAcRNA was hybridised to the Human Genome U133 Plus 2.0 Array (HGU133plus2.0, Affymetrix, SantaClara, CA, USA) and data was initially visualised using MIAMIVICE software. Quality control of microarray data was performed using principal component analysis (PCA) with cross-validation undertaken using Qlucore Omics Explorer 2.3 (Qlucore, Lund, Sweden).

The trophectoderm (TE) and inner cell mass (ICM) of day 6 human embryos were separated by immunosurgically lysing the whole TE (recovering RNA from both mural and polar TE), to leave a relatively pure intact ICM. Eight microarray datasets were obtained, corresponding to 4 genetically paired matched TE and ICM transcriptomes. Frozen robust multiarray averaging (fRMA) (McCall, et al., 2010) was used to define absolute expression by comparison to publically available microarray datasets within R (3.1.2) (Team, 2014). An expression barcode and a z-score of gene expression in comparison to 63331 examples of HGU133plus2.0 was defined for each tissue (McCall, et al., 2014; Zilliox and Irizarry, 2007) and used for unsupervised analysis. For analysis of gene expression specific to each tissue a z-score of 5 was used to call a gene present and a barcode was assigned scoring 1 for presence and 0 for absence of gene expression (McCall, 2015; McCall, et al., 2010; McCall, et al., 2014). All transcriptomic data are available on the Gene Expression Omnibus (GEO) [GSE121982].

### hESC lines

HUES7, HUES3 (kind gift of Kevin Eggan (Cowan, et al., 2004)) and MAN1 (Camarasa, et al., 2010) hESC lines were cultured as previously described (Oldershaw, et al., 2010). Briefly, hESCs (p21-27) were cultured and expanded on Mitomycin C inactivated mouse embryonic fibroblasts (iMEFs) in hESC medium KO-DMEM (Invitrogen, Paisley, UK) with 20% knockout serum replacement (KO-SR, Invitrogen), 8 ng/ml basic fibroblast growth factor (bFGF, Invitrogen), 2 mM L-glutamine, 1% NEAA (both from Cambrex, Lonza Wokingham, UK), and 0.1 mM ß-mercaptoethanol (Sigma-Aldrich, Dorset, UK). For feeder-free culture, cells were lifted from the iMEF layers with TrypLE (Thermo Fisher, Loughborough, UK), and plated onto fibronectin-coated (Millipore) tissue culture flasks with StemPro (Thermo Fisher, Loughborough, UK) feeder-free medium. After 3 passages 100 hESC cells were isolated from each line (assessed separately as > 85% Oct4 positive), lysed and subjected to polyA-PCR amplification, hybridisation to the microarray chip and analysis as described above.

### Analysis of differential gene expression

Principal component analysis was performed to provide further quality control using cross-validation (Qlucore Omics Explorer [QoE] 2.3). Partial least square discriminant analysis (PLSDA) was used to assess the Euclidean distance between the unsupervised transcriptomic samples using the MixOmics package for R (Rohart, et al., 2017).

We analysed published single-cell RNA-Seq data from human epiblast (inner cell mass) and trophectoderm tissue (Blakeley, et al., 2015; Petropoulos, et al., 2016; Yan, et al., 2013). Transcripts per million (TPM) expression values were visualised in QoE and outliers were removed.

### Similarity Network Fusion

Gene probe set similarity network fusion (SNF) (Wang, et al., 2014) was performed on the fRMA derived data as an independent test for similarity, using the *SNFTool* R-package. Euclidean distances were calculated between gene probe sets for each hESC line as well as TE and ICM. Using a non-linear network method based on nearest neighbours, any two of the Euclidean distance matrices could be combined over 20 iterations to produce a final network which accurately describes the relationship between gene probe sets across both initial sets. This method was used to combine each hESC line with TE or ICM gene expression data. The fused data was subjected to spectral clustering to identify groups of gene probe sets with similar patterns of expression across the hESC and TE or ICM samples. This data was presented as a heatmap.

### Hypernetwork assessment of transcriptomic associations

Hypernetworks were generated to understand the relationships between genes which distinguish the trophectoderm and inner cell mass (Stirparo, et al., 2018). Correlations between these transcripts and the rest of the transcriptome were calculated in R (v3.4.4) and the number of shared correlations between pairs of genes was determined (Johnson, 2011). Hierarchical clustering was used to separate a central cluster of genes with high inter-correlation from this network of transcriptomic associations.

### Network model construction and comparison

Lists of differentially expressed genes were used to generate interactome network models of protein interactions related to the transcriptomic data in Cytoscape (Su, et al., 2014) by inference using the BioGRID database (Chatr-Aryamontri, et al., 2015).

The Cytoscape plugin Moduland (Kovacs, et al., 2010; Szalay-Beko, et al., 2012) was applied to identify overlapping modules, an approach that models complex modular architecture within the human interactome (Chang, et al., 2013) by accounting for non-discrete nature of network modules (Kovacs, et al., 2010). Modular hierarchy was determined using a centrality score and further assessed using hierarchical network layouts (summarising the underlying network topology). The overlap between the central module cores (metanode of the ten most central elements) was determined. Community centrality and bridgeness scores were assessed across network models using the Moduland package (Szalay-Beko, et al., 2012). The bridgeness score was used in combination with centrality scores to categorise party and date hubs within the network i.e genes that interact simultaneously or sequentially respectively with neighbours (Komurov and White, 2007; Yu, et al., 2007).

The Network Analyser (Assenov, et al., 2008) Cytoscape plugin was used to calculate associated parameters of network topology. Hierarchical network layouts were used along with centrality scores to assess the hierarchy of network clusters. Significance of the overlap between network elements was calculated using Fisher’s exact test on the sum of each group compared to the expected sum.

The robustness of defined modules is an essential analytical step (Reimand, 2013) and was assessed using permutation analysis in R (version 3.3.2) (RCoreTeam, 2016). Robustness of network module and network topology properties was determined in the ICM and TE interactome network models with 100 permutations of removal of 5, 10, 20, 30, 40 and 50% of the nodes, an approach that has been shown to assess the coherency of network modules (Reimand, 2013). These data were used to assess the stability of network observations.

## Glossary of Network Concepts

Modular Hierarchy: Biological networks form regions of higher connectivity than would be expected by chance, known as modules. Modules represent functionally related elements of a network and their relative influence in a system can be estimated by their centrality.
Metanode: The most central ten connected genes within a module.
Connectivity: The number of links existing between a given node and its neighbours. An increased connectivity is indicative of a gene which is involved in numerous processes.
Community Centrality: A measure of the relative ‘importance’ of a node, characterised by high connectivity or connections between areas of high connectivity.
Bridgeness: A property of a node in a network which sits between two areas of high connectivity, such that if removed, it would cause the separation of a single module into two. These nodes act as ‘bridges’ between modules and an increased bridgeness identifies a node which connects multiple modules.
Party hub: A node with multiple connections which, in a biological system, is thought to represent a gene with many active simultaneous interactions, such as protein complexes. It is characterised by a node which has a reduced bridgeness at a given centrality when compared to a date-hub.
Date hub: A node with multiple connections which, in a biological system, has non-concurrent interactions with other nodes. These are thought to represent transcription factors. It is characterised by a node which has an increased bridgeness at a given centrality when compared to a party-hub.
Similarity Network Fusion: A network approach which uses nearest neighbour relationships to combine datasets and identify regions of similarity within and between them. In the context of this manuscript, coherency between datasets represents genes whose expression patterns are conserved between cells derived from embryonic tissue and human embryonic stem cell lines.

## Acknowledgements

We thank Stuart Pepper at the Christie Genomics unit for Microarray hybridisation. We thank the Medical Research Council UK for funding (grants G0801057 MR/M01 7354/1), and the NIHR clinical research network for support. We would particularly like to thank the research nurses, IVF clinic staff and patients who donated embryos to this research.

## Supplementary Figures

**Supplemental Figure S1.**
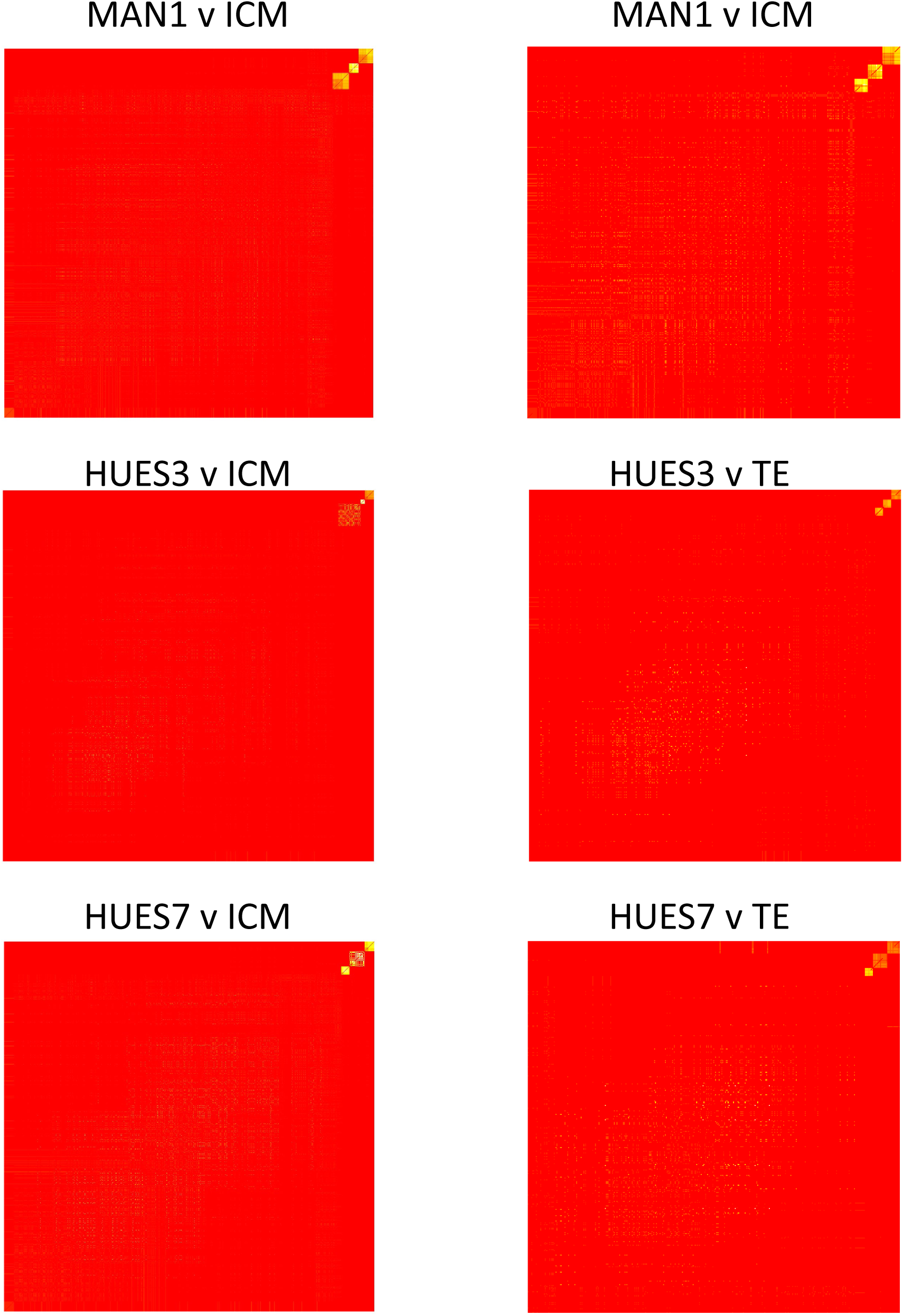
Full similarity network fusion to compare homology between the transcriptome of inner cell mass and trophectoderm and human embryonic stem cell lines. Similarity network fusion matrix showing similarity groups between the uniquely expressed ICM gene probesets from both ICM and the human embryonic stem cell lines (square matrix of gene probesets with leading diagonal showing equivalence mapped to red). Similarity is coloured by intensity from white to yellow, red is dissimilar. The proportion of genes which are similar between a hESC line and either ICM or TE can be determined by the proportion of either axis which contains yellow signal.

**Supplemental Figure 2.**
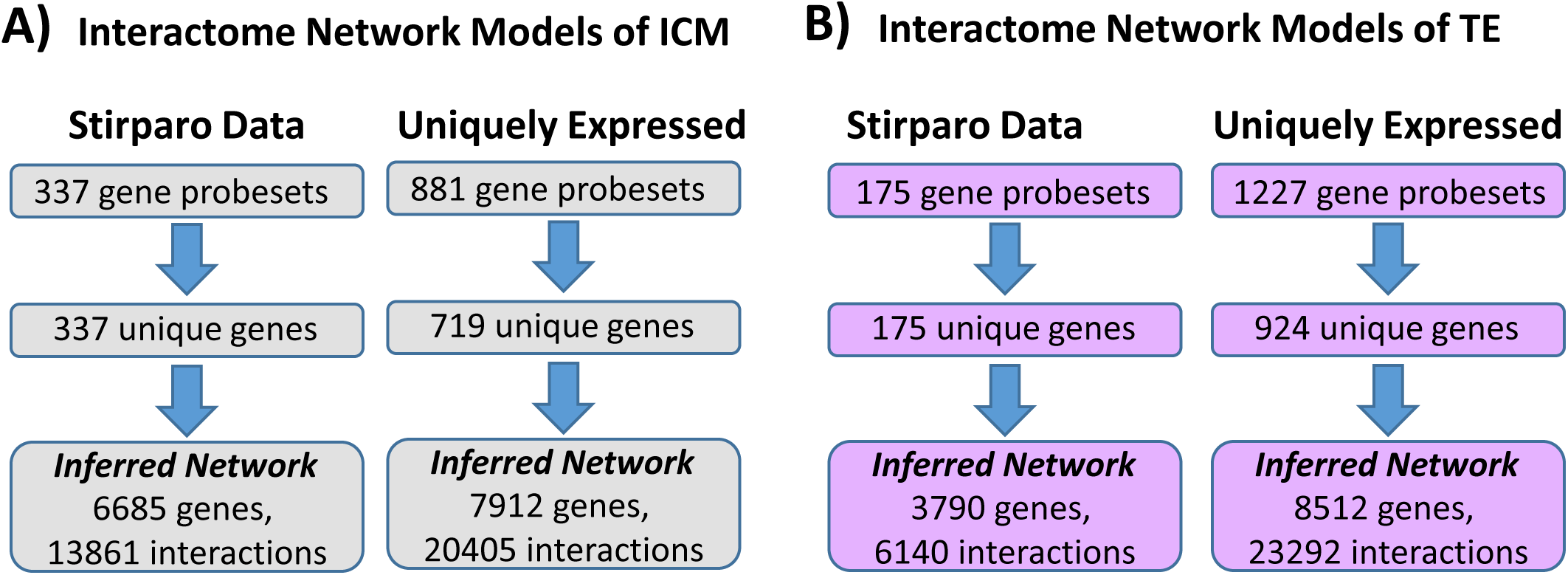
Interactome network models of gene expression in ICM or TE. **A) In Inner Cell Mass (ICM).** Using differentially expressed genes between ICM and TE, unique patterns of transcriptomic expression were defined. Genes with positive expression in ICM (337 from Stirparo meta-analysis) were used to generate an interactome network model for ICM. A second ICM interactome model was generated using the 719 genes (881 gene probesets) uniquely expressed in ICM from our de novo analysis. **B) In Trophectoderm (TE)** The genes differentially expressed between ICM and TE were used (Stirparo datasets from meta-analysis) and genes with positive expression in TE (175) were used to generate an interactome network model. A second TE interactome model was generated using the 924 genes (1227 gene probesets) uniquely expressed in TE from our de novo analysis. These were used to infer interactome network models using the BioGRID database version 3.4.158.

**Supplemental Figure S3.**
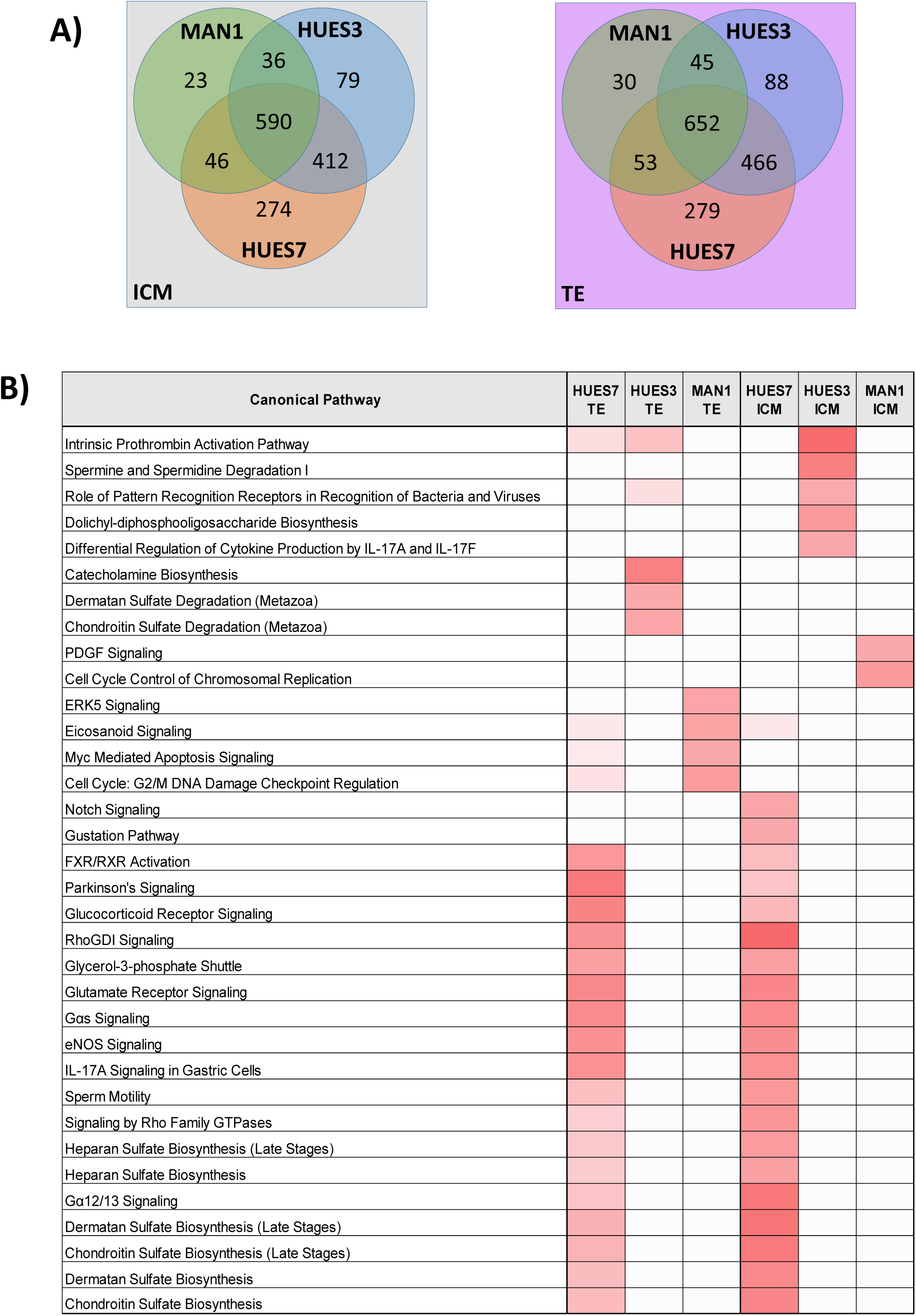
Expressed genes uniquely shared between each human embryonic stem cell line and either the Inner Cell Mass (ICM) or the Trophectoderm (TE). **A)** Overlap of the gene expression (gene probe sets) shared between the human embryonic stem cell lines and ICM or TE. **B)** Biological pathways associated with the gene expression uniquely shared between each human embryonic stem cell line and either ICM or TE. Intensity of red shade is proportional to p-value of right sided Fisher’s Exact test.

**Supplemental Figure S4.**
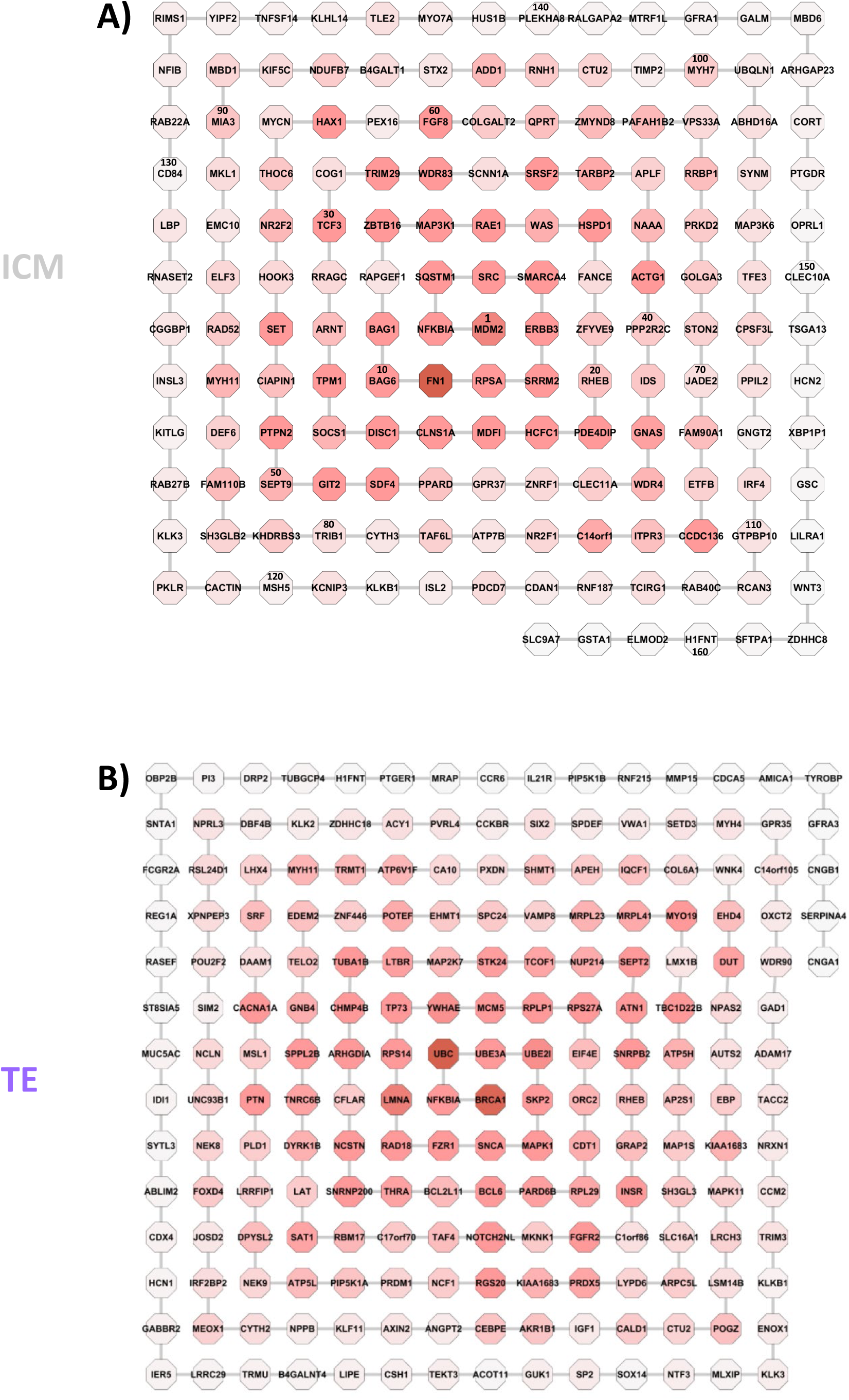
Hierarchy of modules within the interactome network models of ICM and TE based on uniquely expressed genes. **A)** The modules of the ICM and **B)** the TE interactome network represented as octagons named with the most central gene. Modules are arranged in a hierarchy represented as a spiral with numbers defining the position in the hierarchy. Modules are shaded red in relation to connectivity to highlight the relationship between network connectivity and centrality.

**Supplemental Figure 5.**
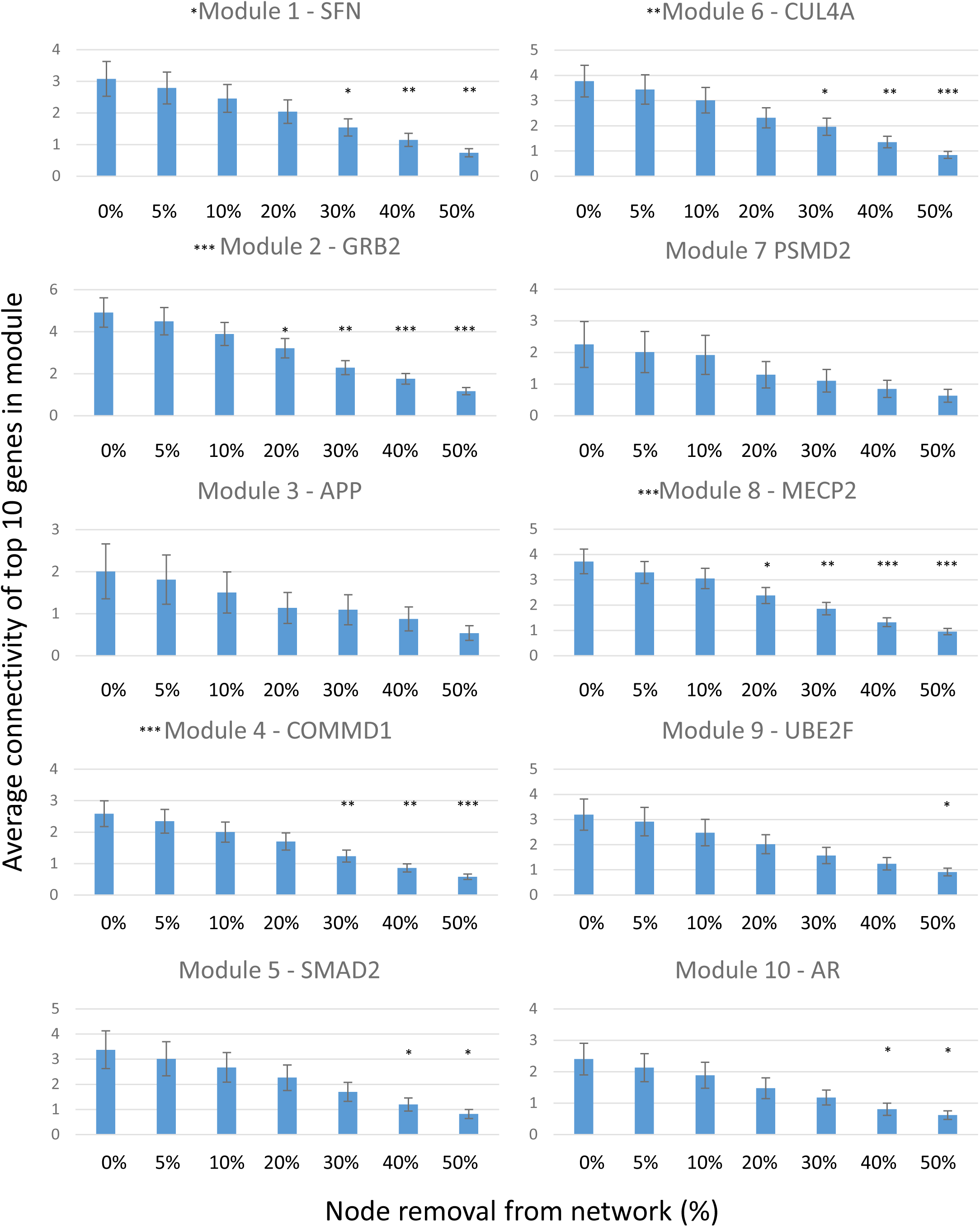
Robustness of 10 most central network modules of an ICM network. Robustness was determined by the mean change in connectivity between the 10 most connected nodes in each network module upon the removal of random nodes from the network. Up to 50% of nodes were removed before recalculating connectivity, iterated 100 times. Significance for each module was determined using ANOVAs whilst between samples t-tests determined significant differences from 0% node loss in each case. Modules whose mean connectivity was not significantly reduced at 20% node removal can be described as robust. [* p < 0.05; ** p < 0.01; *** p < 0.001].

